# Amyloid-beta relates to default and attention network connectivity during working memory retrieval

**DOI:** 10.64898/2025.12.21.695827

**Authors:** Hengda He, Christian Habeck, Yaakov Stern

## Abstract

As one of the earliest pathological events in Alzheimer’s disease, amyloid-beta accumulation has been linked to alterations in large-scale brain networks, especially the default mode network. While most prior work has focused on resting-state functional connectivity or task-evoked activation, how amyloid-beta relates to task-modulated connectivity that directly supports behavior is less explored. Here, we examined the relationship between amyloid-beta burden and task-modulated connectivity during a Letter Sternberg verbal working-memory task in eighty-four cognitively normal older adults from an ongoing longitudinal cohort (age 56-71 years; 41 female). To further understand these connectivity alterations, we tested whether these connections were associated with task performance or with cognitive reserve factors. We found that higher amyloid-beta burden was associated with reduced connectivity specifically during the probe phase of working-memory retrieval. At the regional scale, lower node strength in posterior-medial regions of the default mode network was related to worse task performance and mediated the amyloid-behavior relationship. At the network scale, altered coupling between the default mode and dorsal attention networks was related to task performance but did not mediate the amyloid-behavior relationship. Moreover, physical activity was associated with network-level coupling but not with regional node strength. These findings suggest that amyloid-beta burden disrupts retrieval-related working-memory systems, particularly regions of the default mode network that are vulnerable to amyloid-beta, while coupling between the default mode and dorsal attention networks indexes resilience. Collectively, task-modulated connectivity provides insight into pathways through which early Alzheimer’s disease pathology influences working-memory performance and resilience and may inform targeted lifestyle or neuromodulation interventions for older adults.

## Introduction

Many older adults remain cognitively normal despite accumulating pathological proteins in the brain. However, the buildup of amyloid-beta (Aβ) and tau aggregates^1^ is associated with an increased risk of future cognitive decline^2^, marking the preclinical phase of Alzheimer’s disease (AD)^3^. Among these pathological processes, Aβ deposition is considered one of the earliest initiating events in the disease cascade and can trigger downstream neuropathological changes, including tau propagation, synaptic and network dysfunction, and ultimately cognitive impairment^4,5^. Growing evidence indicates that large-scale functional brain networks, particularly the default mode network (DMN), are vulnerable to such early pathological disruptions, with such alterations potentially linked to cognitive decline^6–8^. However, it is not fully understood how early AD pathology, such as Aβ, interacts with functional brain networks during task performance. Furthermore, individual differences in cognitive reserve factors, including education, IQ, and physical activity, may also influence brain network alterations or adaptations in the presence of Aβ pathology^9–12^, but the functional pathways underlying these resilience mechanisms are not fully established. We examined how Aβ burden relates to task-modulated functional connectivity during a working memory task and tested how these Aβ-associated connectivity changes relate to cognitive performance and cognitive reserve factors in cognitively normal older adults.

Recent studies have demonstrated that early Aβ deposition preferentially accumulates in and affects the DMN, especially its highly connected hub regions, representing regional vulnerability to early AD pathology^13–15^. However, most existing neuroimaging findings are based on resting-state functional MRI (fMRI) and correlation-based connectivity measures, which limit interpretation regarding whether altered connectivity reflects pathological vulnerability or adaptive compensation^16^. Task-modulated connectivity derived from task-based fMRI captures dynamic, context-dependent interactions among brain regions, which are distinct from correlation-based estimates of spontaneous fluctuations^17^. Recent evidence using task-based connectivity approaches suggests that Aβ-related alterations in network connectivity may influence downstream neuropathological processes, including tau accumulation^18,19^. In addition to Aβ-related hub region vulnerability, the functional activity of the DMN and dorsal attention network (DAN) is implicated in cognitive reserve mechanisms, where these networks may adaptively compensate for the impact of pathology or neurodegeneration on cognition^20,21^.

Therefore, alterations in these large-scale networks during aging and early AD may represent both local vulnerability to pathology and network-level compensatory adaptation that supports cognitive function. Accordingly, brain-behavior relationships related to Aβ burden may be more tightly linked to regional hub connectivity, whereas associations with cognitive reserve factors may emerge more prominently at the network level. We therefore hypothesized that Aβ-related disruptions in DMN hub connectivity are associated with poorer task performance, while DMN couplings with the DAN could reflect adaptive resilience.

Working memory (WM) refers to the cognitive systems responsible for the temporary maintenance and manipulation of information, which are critical for supporting higher-order functions such as reasoning, decision-making, and goal-directed behavior^22^. WM is known to decline with aging and AD^23^, and is disrupted in the early stages of AD^24^. In the current study, participants performed a Letter Sternberg (LTS) verbal WM task during fMRI, which included three task phases: encoding, retention, and probe/retrieval, as well as three memory-load conditions. Previous studies have shown that Aβ pathology influences WM-related functional activity. For example, Oh et al. reported Aβ-associated load-dependent hyperactivity in the frontoparietal network (FPN)^25^ during a working memory task, and Kennedy et al. found that Aβ burden interacts with WM-related brain function in a nonlinear fashion, affecting both task performance and executive control^26^. More recently, Argiris et al. demonstrated that phase-specific patterns of WM functional activation across encoding, retention, and probe may represent a brain maintenance mechanism in the presence of neurodegeneration^27^. These findings highlight the importance of examining WM task phases separately. However, it remains unclear which specific phase of WM is most affected by Aβ pathology and how such effects contribute to cognitive performance. Furthermore, examining Aβ-related alterations in connectivity and their relationships to task performance, as well as reserve and resilience factors, has important implications for early intervention design, such as identifying optimal network targets for transcranial magnetic stimulation (TMS) to enhance cognitive functioning^28,29^, and establishing connectivity-based markers to monitor the progression and efficacy of physical activity interventions^30,31^.

In this study, we first examined task-modulated functional connectivity during a WM task using a whole-brain psychophysiological interaction (PPI) analysis^32^. We then investigated how Aβ burden, as estimated from PET imaging, was related to task-modulated connectivity across different phases of the task. To assess whether Aβ-related connectivity alterations were associated with task performance or with reserve and resilience factors, we quantified WM efficiency, calculated as accuracy divided by response time (RT), and assessed reserve and resilience factors with IQ, education, and physical activity. To test whether Aβ-related effects differ across spatial scales, we examined task-modulated connectivity at both the regional and large-scale network levels. We hypothesized that higher Aβ burden would be related to reduced connectivity of DMN hub regions, which in turn would be associated with poorer task performance. Furthermore, we hypothesized that higher network-level connectivity, particularly between the DMN and DAN, would be related to higher resilience factors, reflecting adaptive network reconfiguration in the presence of Aβ pathology. Together, this study provides insight into how Aβ burden disrupts WM-related connectivity and identifies potential resilience pathways that may inform designs to promote cognitive reserve and resilience.

## Methods

### Participants and study design

Participants were drawn from the ongoing Cognitive Reserve/Reference Ability Neural Network (CogRes/RANN) longitudinal study^33^. Recruitment was conducted through random market mailings in the New York metropolitan area. Individuals were screened to ensure they were right-handed, native English speakers with no major neurological or psychiatric disorders, history of head injury, significant medical illness, or hearing/vision impairment that could interfere with MRI acquisition or cognitive testing. All participants were cognitively normal at data acquisition, and dementia or mild cognitive impairment (MCI) was excluded using the Dementia Rating Scale (DRS)^34^, with a minimum inclusion score of 130. The experimental design of our study and the recruitment process were approved by the Internal Review Board of the College of Physicians and Surgeons of Columbia University. All participants have provided informed consent to participate in the study, and written consent was obtained from the participants. For the present analysis, we included subjects who had completed baseline structural and task-based functional MRI and amyloid PET imaging and had available demographic data.

### Letter Sternberg working-memory task paradigm and behavioral measure

As illustrated in Figure 1A, during fMRI acquisition, participants performed an LTS verbal WM task with three memory-load conditions (one, three, or six letters). Each trial began with the presentation of an upper-case letter set for 3 sec (encoding phase), followed by a 7 sec of blank screen (retention phase), and ended with a lower-case probe letter displayed for 3 sec (probe/retrieval phase). Participants indicated by button press whether the probe letter was included in the original set. Each participant completed 90 trials across three fMRI runs, with a jittered inter-trial interval (gamma distribution; k = 1.3, θ = 2) to randomize trial onset. Participants were instructed to respond as accurately and quickly as possible and were trained on the task before scanning. Responses were collected with a fiber-optic button box by button press. The primary behavioral measure of LTS working-memory tasks was WM efficiency. This task performance measure was calculated as accuracy divided by mean RT and then averaged across three memory loads. Mean accuracy and mean RT were also examined separately.

**Figure 1.**
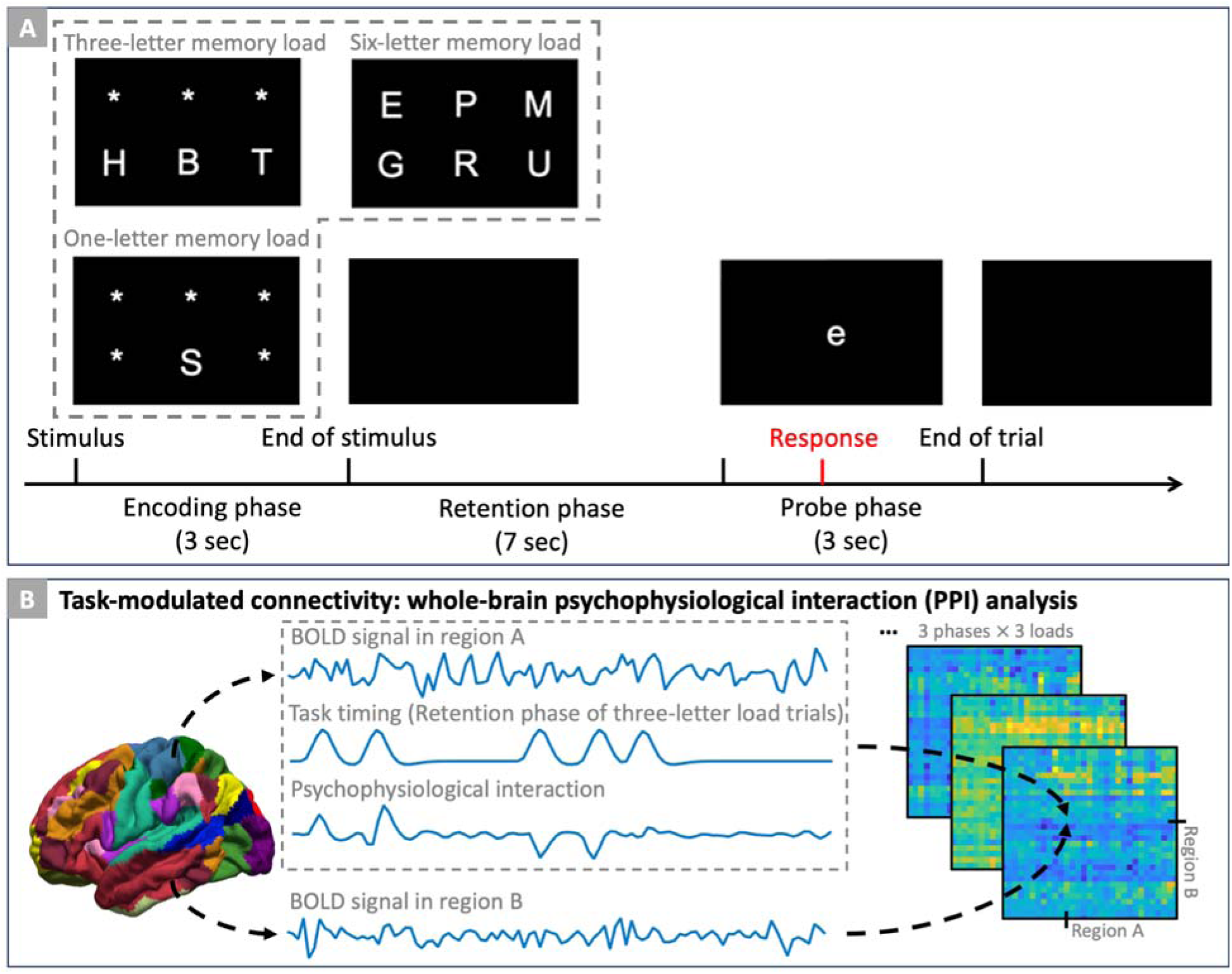
Functional MRI task paradigm and task-modulated functional connectivity analysis. (A) Letter Sternberg verbal working-memory task paradigm. Each trial started with a presentation of an upper-case letter set for 3 sec (encoding phase), followed by a 7 sec blank screen (retention phase), and a 3 sec presentation of a lower-case probe letter (retrieval phase). Participants were instructed to press the button indicating whether the probe letter was in the letter set during the encoding phase. Memory load (one, three, or six letters for encoding) was randomized across trials. Task performance was quantified as efficiency (accuracy divided by median response time). (B) Task-modulated psychophysiological interaction (PPI) functional connectivity analysis. Whole-brain PPI models were used to estimate connectivity between regions of interest from the Schaefer atlas. For each participant, nine connectivity matrices (3 phases × 3 loads) were estimated by modeling the interaction between regional BOLD signals and working-memory task timing, controlling for task-evoked activation and nuisance covariates

### MRI and PET data acquisition

All MRI data were collected on a 3T Philips Achieva Magnet scanner. Anatomical high-resolution T1-weighted images were acquired using a magnetization-prepared rapid gradient echo (MPRAGE) sequence (TR/TE = 6.6/3.0 ms; flip angle = 8°; field of view (FOV) = 256 × 256 mm; matrix = 256 × 256; voxel size = 1 mm isotropic; 165 axial slices). A neuroradiologist confirmed the absence of clinically significant abnormalities in all participants. LTS working-memory task-based fMRI scans were collected with a T2*-weighted echo-planar imaging (EPI) sequence (TR/TE = 2000/20 ms; flip angle = 72°; FOV = 224 × 224 mm; matrix = 112 × 112; slice thickness = 3 mm; 36 axial slices, 314 volumes). Three runs of approximately 10.5 min each were collected. The first three volumes of each run were discarded to allow for magnetization equilibrium.

Amyloid PET imaging used [18F]Florbetaben (donated by Piramal Pharma, Inc.) tracer on a Siemens Biograph 64 mCT/PET scanner (3D dynamic acquisition, four 5-min frames, 20 min total). Data acquisition began 50 min after a 10 mCi dose bolus injection of 18F-florbetaben. A structural CT scan (0.58 × 0.58 mm in-plane resolution; 3 mm slice thickness) was also acquired for attenuation correction purposes.

### MRI and PET data processing

Functional MRI preprocessing was performed using FSL (v6.0.4) and custom Python code. The pipeline included motion correction (MCFLIRT^35^) with the first volume as the reference volume, slice-timing correction with *slicetimer*^36^, temporal high-pass filtering (> 0.01 Hz) to remove scanner drift, and spatial smoothing with a 5 mm full-width at half-maximum Gaussian kernel.

Structural T1-weighted images were processed with FreeSurfer (v5.1) for tissue segmentation and cortical surface reconstruction. Cortical parcellation was performed with the Schaefer atlas^37^ (200 cortical parcels with 17 networks version). Specifically, the surface-space Schaefer parcellation was registered to everyone’s cortical surface space and projected to the native EPI volumetric space. Tissue segmentation and registrations were visually inspected for quality control.

Amyloid PET images were processed with an in-house developed pipeline^38^. Briefly, PET frame images were registered to the first frame image with rigid-body transformation, averaged to generate a static PET image. Then this image was co-registered to each participant’s CT image to create a PET-CT composite image. This image was later registered to each participant’s structural T1-weighted brain image. The Schaefer parcellation regions of interest (ROIs) and the cerebellar gray matter mask (FreeSurfer segmentation) were transformed into the PET image space. Lastly, standardized uptake value ratios (SUVRs) were calculated for each ROI by normalizing regional tracer uptake to the cerebellar gray-matter reference. Aβ burden was calculated as the mean SUVR across cortical regions previously identified linked to longitudinal cognitive decline in global cognition (defined as Aβ signature regions; more details are in a previous study^39^; regions include temporo-occipital, orbitofrontal, and ventrolateral prefrontal cortex). A global SUVR was also computed across the entire cortex.

### Task-modulated functional connectivity

To examine the task-modulated functional connectivity during the working-memory process, we performed a whole-brain PPI analysis (Figure 1B). For each participant, we first extracted the blood oxygenation level dependent (BOLD) time series of every cortical region defined by the Schaefer parcellation. Task-modulated connectivity was then estimated using multiple-regression, where the BOLD signal of each region (ROI A) was modeled as a linear combination of: 1) the BOLD signal of a second region (ROI B); 2) task timing regressors representing the onsets and durations of the three working-memory task phases (encoding, retention, probe) at each of the three memory loads (one-, three-, and six-letter sets); 3) the PPI interaction terms formed by multiplying the deconvolved BOLD signal of ROI B with each of the nine phase-by-load task boxcar function, and re-convolving with the canonical hemodynamic response function (HRF); 4) controlling for nuisance covariates (motion parameters, large motion volumes, mean BOLD signals from white matter and lateral ventricles, and constant terms for each run). The inclusion of task timing regressors ensured that task-evoked mean activation was controlled, thereby isolating task-modulated connectivity effects from task co-activation. To further control potential false positive results^40^, FMRIB’s linear optimal basis set (FLOBS)^41^ was used to model task timing regressors. Specifically, a boxcar function representing the timing of task trials for each phase and load was convolved with the FLOBS basis set, yielding a total of twenty-seven task timing regressors. For each subject, the PPI model resulted in nine task-modulated connectivity matrices (three phases by three loads). Beta coefficients of the interaction terms represented the strength of task-modulated connectivity between each pair of regions. Lastly, we also calculated positive and negative node strength, as the sum of all positive or negative edge weights associated with each ROI.

### Statistical Analysis

All analyses controlled for age and sex. We first tested the relationship between Aβ burden and LTS working-memory task performance. Average Aβ SUVR in the signature regions were correlated with working-memory task efficiency performance, RT, and accuracy, using partial Spearman correlation and controlling for covariates.

To investigate whether task-modulated connectivity exhibits phase-dependent or load-dependent effects, repeated-measures ANOVA tests were used on edgewise PPI connectivity, with false-discovery rate (FDR) correction at q < 0.05 across edges. Group-level task-modulated connectivity for each task phase was computed using a one-sample t-test (FDR-corrected p < 0.05) with connectivity matrices averaged across the loads.

Next, the task-modulated connectivity was related to amyloid burden. For each phase, partial Spearman correlations were used to assess associations between amyloid SUVR and PPI connectivity averaged across memory loads. Significant effects were identified with FDR correction (q < 0.05) across edges between pairs of ROIs. Then, positive and negative node strength was also related to amyloid burden, and significant relationships were identified with FDR correction. Lastly, node-level connectivity was summarized into network-level connectivity based on the network label of each node from the Schaefer atlas network assignment. Network-level connectivity of each phase was correlated with the amyloid burden, and significant edges were extracted with FDR-corrected p < 0.05.

To assess the associations between amyloid-related connectivity and behavioral outcomes, we implemented a 20-fold cross-validation (CV) framework with 500 random repetitions. This repeated K-fold strategy was chosen because it provides more stable and less biased estimates than leave-one-out approaches, particularly in moderate sample sizes^42^. Specifically, in each repetition, participants were randomly partitioned into 20 folds. For each fold, a linear regression model was trained on 19 folds using the amyloid-related node- or network-level averaged connectivity measures as a predictor of either task performance or resilience factors, with age and sex included as covariates. The trained model was then applied to the held-out fold to generate out-of-sample predictions. This process was iterated until every fold was used as the test set once, resulting in predicted values for all participants. The entire 20-fold CV procedure was repeated 500 times with different random partitions to minimize variability due to data splits. Predictive performance was quantified as the median Pearson correlation coefficient between predicted and observed outcomes across repetitions. To assess the significance of the cross-validated correlation, permutation testing (1000 iterations) was used by randomly shuffling the dependent variable to generate a null distribution. The p-value was computed as the proportion of permutations that were greater than or equal to the observed cross-validated correlation. CV frameworks with different selection of fold number K were also tested, and results were reported in the Supplementary Text.

For task-modulated connectivity measures that demonstrated a significant relationship to task performance, we tested whether these connectivity measures mediated the influence of amyloid burden on working-memory task performance. A bootstrapped mediation model was used with 5000 resamples, and we reported 95% confidence intervals and two-tailed p-values for both the indirect mediated and direct effects.

## Results

### Study participants

Eighty-four participants from the CogRes/RANN study with available datasets were included in the analyses (age range 56-71 years; mean ± standard deviation (SD) = 65.5 ± 3.4; 41 females), with education ranging from 12 to 20 years (mean ± SD = 16.3 ± 2.2). All participants performed the LTS working-memory task well, with mean WM efficiency of 0.87 (SD = 0.20, range from 0.54 to 1.53), mean RT of 1.19 sec (SD = 0.24, range from 0.66 to 1.68), and mean accuracy of 0.95 (SD = 0.04, range from 0.78 to 1.00). The mean global amyloid SUVR among the participants was 1.16 (SD = 0.09) with three out of eighty-four participants classified as amyloid-positive using a cutoff of 1.34 as defined from a previous study^43^.

### Association of amyloid burden and working memory task performance

We first examined whether amyloid burden in signature regions was related to working-memory task performance. The primary behavioral outcome was task efficiency, calculated as accuracy divided by mean RT and averaged across memory loads. Higher signature-region amyloid SUVR was significantly associated with poorer task performance (*r* = -0.3237, *p* < 0.0032, controlling for age and sex). We also tested global SUVR and individual performance components. A similar negative association was observed for global amyloid SUVR (*r* = -0.2348, *p* < 0.0349). Higher amyloid burden was further related to slower response times (signature: *r* = 0.3219, *p* < 0.0034; global: *r* = 0.2310, *p* < 0.0380) and lower accuracy (signature: *r* = -0.2773, *p* < 0.0122; global: *r* = -0.1996, *p* = 0.0740).

### Task-modulated connectivity supporting working memory

We next examined task-modulated connectivity using PPI analysis. For each subject, nine connectivity matrices were derived, reflecting the three memory loads (one-, three-, and six-letter sets) across the three task phases (encoding, retention, probe). A repeated-measures ANOVA revealed a significant main effect of phase on task-modulated connectivity (FDR-corrected *p* < 0.05; Figure 2A), but no significant effect of memory load. Accordingly, connectivity measures were averaged across memory load conditions for subsequent analyses. We then identified significant group-level connectivity for each phase: 1) during encoding, the strongest positive edge was observed between RH-SalVentAttnA and RH-DefaultA networks, and the strongest negative edge between RH-DorsAttnB and LH-DorsAttnB networks (Figure 2B); 2) during retention, the strongest positive edge was between RH-SalVentAttnA and LH-ContB networks, and the strongest negative edge between RH-SomMotA and LH-SomMotA networks (Figure 2C); and 3) during the probe phase, connectivity was most positive between LH-DefaultA and LH-VisCent networks and most negative between RH-SomMotA and LH-SomMotA networks (Figure 2D).

**Figure 2.**
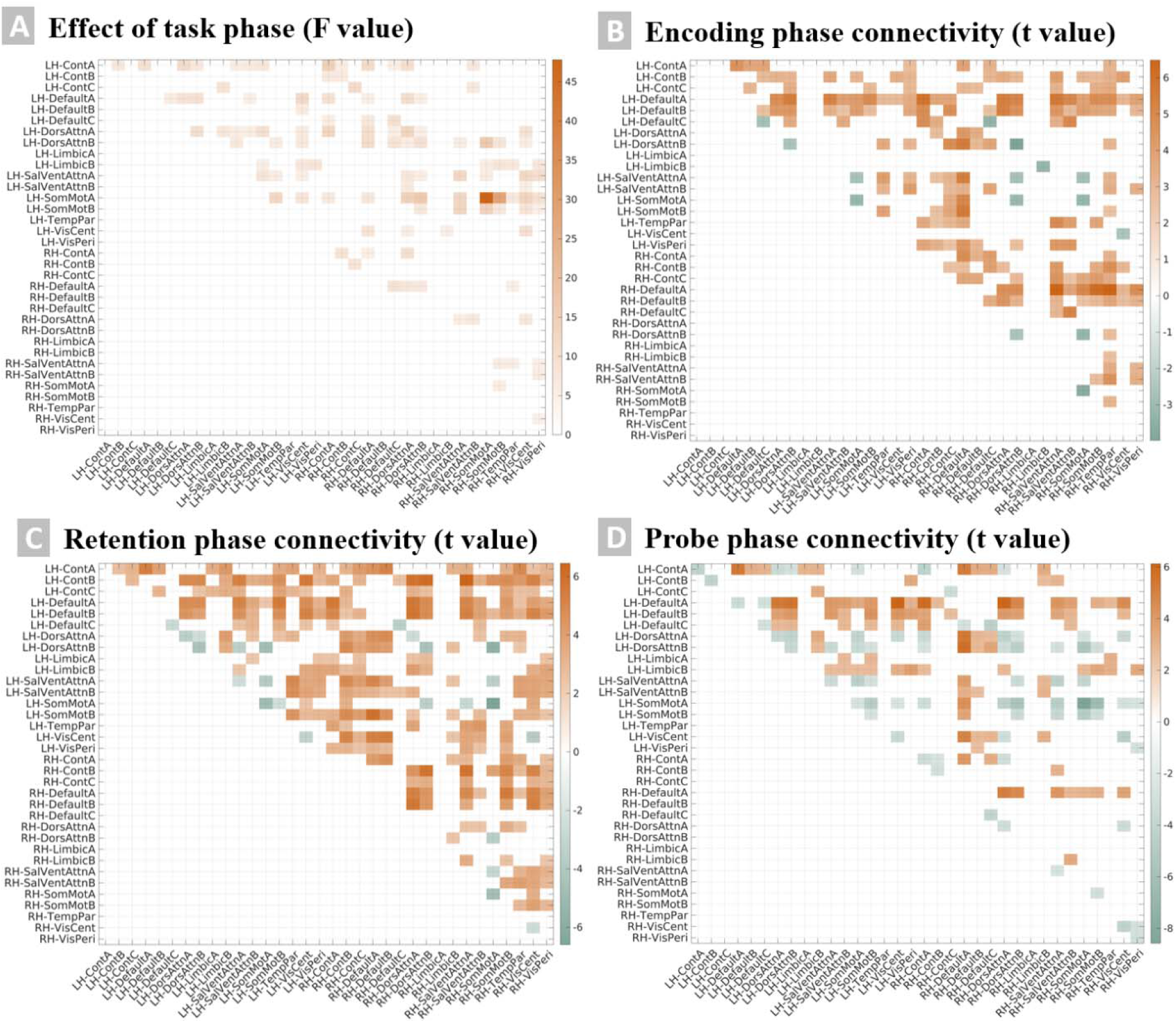
Task-modulated psychophysiological interaction (PPI) connectivity during the Letter Sternberg working-memory task. (A) Repeated-measures ANOVA showing the main effect of task phase (encoding, retention, probe) on PPI connectivity across all memory loads (F values, FDR-corrected p < 0.05). (B–D) Significant group-level task-modulated connectivity (t values) identified for the encoding (B), retention (C), and probe (D) phases. Positive t values (orange) indicate increased connectivity during the task relative to baseline, whereas negative t values (green) indicate decreased connectivity. Connectivity matrices represent pairwise connections across left (LH) and right (RH) hemisphere cortical networks based on the Schaefer atlas brain parcellation.

### Amyloid burden is associated with altered probe-phase connectivity

We tested whether amyloid SUVR in signature regions was related to task-modulated connectivity across phases of the working-memory task, with connectivity averaged across memory loads. No significant associations were observed for encoding or retention phases. In the probe phase, however, higher amyloid burden was associated with reduced connectivity at both the region and network levels (Figure 3). At the node level (Figure 3B), higher amyloid was linked to reduced connectivity between the right retrosplenial area (Rsp) and the right somatomotor area, as well as lower positive node strength in the bilateral parahippocampal cortex (PHC), the right retrosplenial area, and the right inferior parietal lobule. At the network level (Figure 3C), higher amyloid was associated with reduced connectivity between the DMN and both the DAN and somatomotor network (specifically, between the bilateral DMN and the bilateral DAN, and between the right DMN and the right somatomotor network). Analyses using global amyloid SUVR showed convergent but weaker effects (Table SI and Table SII). For comparison purposes, relationships between amyloid burden and resting-state functional connectivity were also tested, and no significant relationship was observed with FDR-corrected p < 0.05 (more details in Supplementary Text and Figure S1).

**Figure 3.**
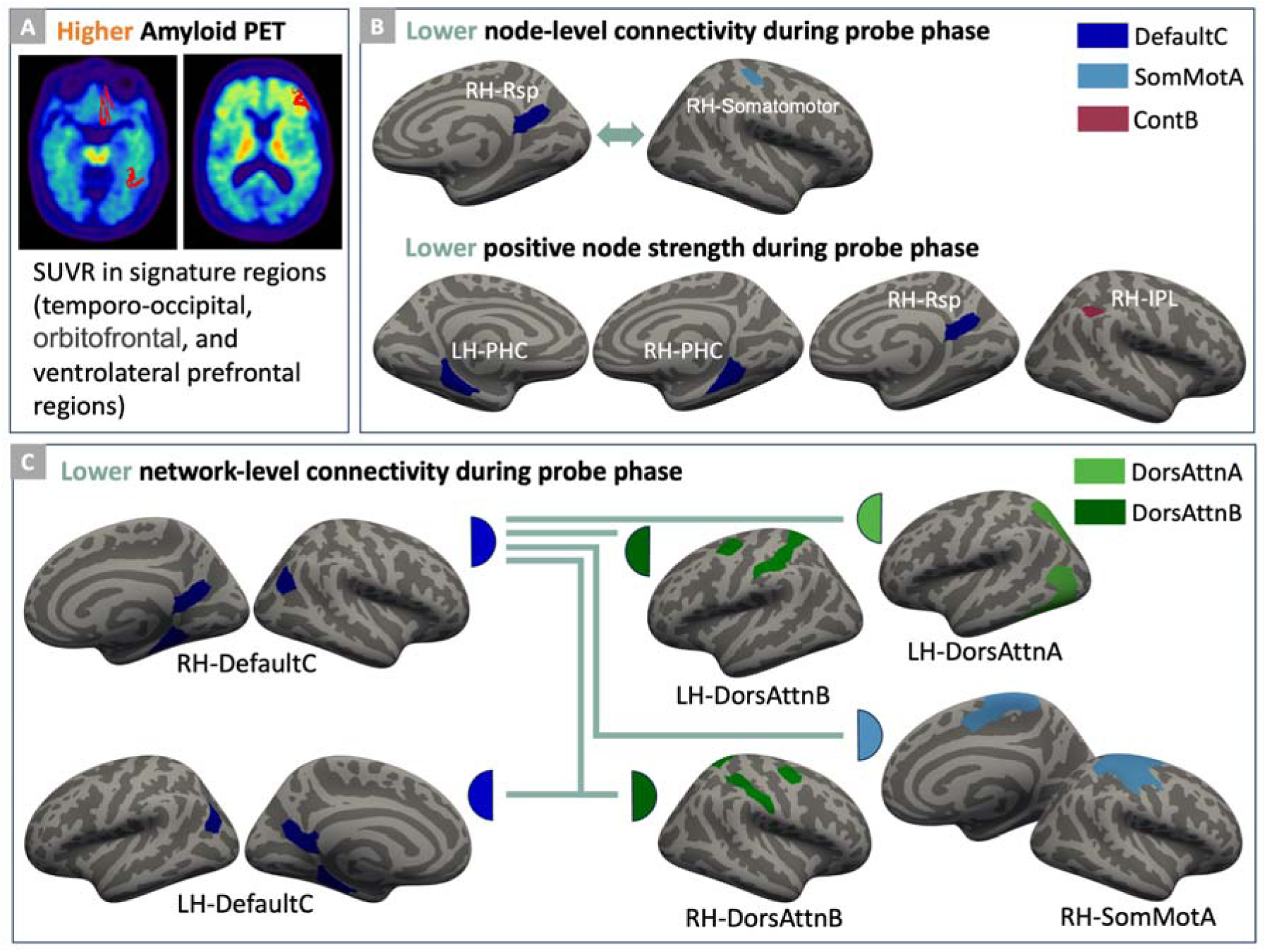
Amyloid-associated changes in probe-phase task-modulated connectivity. (A) PET images were used to capture amyloid burden within the predefined signature regions. (B) Node-level amyloid-connectivity associations during the probe phase. Higher amyloid-beta burden was linked to lower connectivity between the right retrosplenial area (RH-Rsp) and the right somatomotor area (RH-Somatomotor), and to lower positive node strength in the bilateral parahippocampal cortex (LH-PHC, RH-PHC), RH-Rsp and right inferior parietal lobule (RH-IPL). (C) Network-level amyloid-connectivity associations during the probe phase. Higher amyloid-beta was associated with lower connectivity between the default mode network and both the dorsal attention and the somatomotor networks.

### Regional node strength mediates the association between amyloid-beta burden and task performance

We next tested whether regional node strength was related to task performance. Positive node strength values from the regions identified in the previous analysis (Figure 3B) were averaged and included in a mediation model. Higher averaged node strength was positively associated with better task performance (*r* = 0.3899, *p* < 3.2015e-04, Spearman correlation controlling age and sex). Mediation analysis indicated a significant indirect effect of amyloid burden on task performance through regional node strength (mediation effect = -0.1297, 95% confidence interval = [-0.2491, -0.0227], p < 0.0240), while the direct effect of amyloid on task performance was not significant (c′ = -0.1910, 95% confidence internal = [-0.4319, 0.0182], p = 0.0728) (Figure 4A). Analyses using global amyloid SUVR showed a similar mediation effect (mediation effect p < 0.0064; Supplementary Text). Amyloid-associated connectivity between the right Rsp and the right somatomotor area (Figure 3B) was not significantly related to task performance (*r* = 0.2097, *p* = 0.0602).

**Figure 4.**
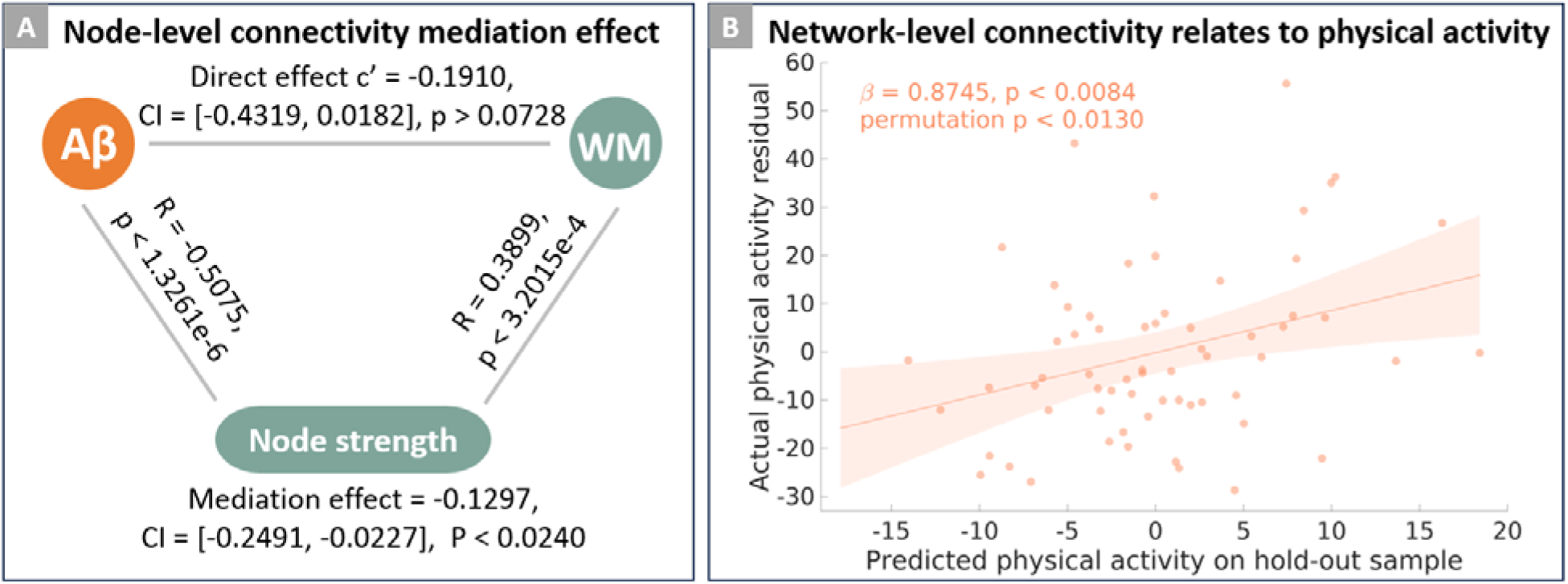
The relationships between working memory (WM) task performance, physical activity lifestyle resilience factor, and amyloid-beta (Aβ) related node-level and network-level connectivity. (A) Mediation analysis demonstrated a significant mediation effect of node-level connectivity during WM retrieval in the relationship between Aβ and WM task performance. (B) Cross-validated prediction of physical activity from Aβ-related network-level connectivity. Connectivity between default mode and dorsal attention networks significantly predicted physical activity. The shaded area indicates the 95% confidence interval of the regression line.

We also tested the relationship between regional node strength and task performance using cross-validation. Averaged node strength values from the identified regions were used as an independent variable in a 20-fold cross-validation framework with 500 repeats and permutation testing. Regional node strength was significantly related to task performance (permutation *p* < 0.0020), with results consistent across different cross-validation folds (Supplementary Text).

Similarly, the averaged network connectivity from the identified edges (Figure 3C) was also significantly related to task performance (cross-validation permutation *p* < 0.0150). However, averaged network connectivity did not significantly mediate the effect of amyloid burden on task performance (mediation effect = -0.0656, 95% confidence interval = [-0.1760, 0.0408], p = 0.1772).

### Amyloid-related network-level connectivity is associated with physical activity

We examined whether amyloid-related network-level connectivity was associated with proxies of cognitive reserve and resilience, including education, IQ and physical activity. Cross-validation analyses showed no significant associations between averaged network-level connectivity and education (permutation p = 0.6903) or IQ (permutation p = 0.3836). Amyloid-associated network-level connectivity was significantly related to physical activity (permutation *p* < 0.0130; Figure 4B). However, network-level connectivity did not significantly moderate the effect of amyloid burden on task performance (details in Supplementary Text). Amyloid-related regional node strength measure was not significantly related to physical activity (permutation *p* = 0.2068).

Next, in a subgroup of participants with follow-up data available, we further tested the relationship between physical activity and the DMN-DAN network connectivity, based on the amyloid-related connectivity edges (Figure 3C). Longitudinal analyses showed that baseline physical activity was significantly associated with changes in DMN-DAN connectivity over time (RH-DefaultC averaged connectivity with LH-DorsAttnA, LH-DorsAttnB, and RH-DorsAttnB: r = 0.4573, p < 0.0490; LH-DefaultC and RH-DorsAttnB: *r* = 0.5272, *p* < 0.0204; *N* = 22; Pearson correlation controlling age, sex, and the corresponding baseline connectivity).

## Discussion

In this study, we investigated how Aβ burden relates to task-modulated connectivity during distinct phases of a working memory task. We found that higher Aβ was specifically associated with reduced connectivity between the default mode and dorsal attention networks during the probe (WM retrieval) phase, but not during encoding or retention phase. At the regional level, Aβ burden was linked to lower node strength in posterior-medial (PM) network regions, including the PHC and Rsp, and this mediated the relationship between Aβ and working-memory task performance. Furthermore, we observed a significant relationship between Aβ-related network connectivity and physical activity, indicating a potential pathway through which lifestyle factors may enhance resilience in the face of early AD pathology. Together, these findings identify a phase-specific task connectivity mechanism through which Aβ disrupts working-memory processes and suggest that lifestyle factors such as physical activity may help promote resilience.

Working memory refers to cognitive processes responsible for temporal maintenance and manipulation of information, involving both attentional and executive control^22^. Working memory impairment is present in the very early stages of AD and has been widely studied in normal aging and across the AD continuum^23,24^. In this study, we found that Aβ burden was specifically associated with altered task-modulated connectivity during the probe phase of a working memory task, but not during encoding or retention. The probe phase of working memory involves complex functional subprocesses, including stimulus identification and recognition, working memory readout, decision making, and motor response, where the participants are required to retrieve information from working memory system and allow it to acquire control of behavior^44,45^. The selective association of Aβ with probe-phase connectivity may reflect amyloid-related disruptions in the neural mechanisms supporting working memory retrieval. Similar task phase-specific findings have been reported for episodic memory retrieval, where Aβ related to decreased connectivity only for retrieval phase not encoding and maintenance phase^46^. Our results suggest a potential shared Aβ vulnerability of memory retrieval during both episodic memory and WM, likely impacting through the episodic buffer component of working memory^47^.

The PHC and Rsp are considered the PM network, which plays an important role in human memory, especially for recollection-based memories, memory for spatial and episodic context, and scene perception^48^. We found that higher amyloid was associated with reduced connectivity among the PHC, Rsp, and inferior parietal lobule. This aligns with previous reports showing that Aβ deposition preferentially affects the PM networks^49–51^. At the network-level, we found that higher amyloid was linked to reduced connectivity between the DMN and both the DAN and somatomotor (SMN) networks. This aligns with prior work showing that Aβ pathology disrupts DMN^8,52^. Additionally, a loss of resting-state functional connectivity between the DMN with the DAN and SMN has been related to progression of AD^53–55^. Together, our results extend these findings by demonstrating that Aβ-related disconnections emerge not only at rest but also during working memory task execution and specifically during the retrieval phase.

At the behavioral level, our cross-validation analyses showed that amyloid-disrupted regional node strength and network-level connectivity can both significantly predict task performance. However, mediation analyses revealed that only regional node strength significantly mediated the relationship between Aβ burden and task performance. These results support previous studies showing that Aβ or neurodegenerative pathology interacts with brain connectivity to influence cognitive performance^26,46,50,56–58^. While our sample in the current study is dominated by amyloid-negative cognitively normal individuals, previous studies suggest that there exists a nonlinear relationship between Aβ and brain function^26,58^, which likely varies across the stages of the AD continuum^16,50^. For example, increased DMN-DAN connectivity was related to compensation in cognitively normal groups, whereas this hyperconnectivity also represented worse memory in an MCI group^56^. Future work is needed to examine the relationships between Aβ burden, task connectivity, and task performance at different stages of the AD continuum.

Aβ deposition has been consistently linked to alterations in brain activation and connectivity. However, it remains challenging to interpret whether changes in connectivity represent vulnerability to Aβ pathology, adaptive resilience, or both^59^. However, disentangling these effects are critical for both biomarker development and intervention design^60^. For example, some studies proposed altered connectivity as compensatory processes^61,62^, whereas others view such alterations as detrimental network breakdown^1,8^. To address this, we examined whether Aβ-associated connectivity was related to resilience factors. We found that the altered network-level connectivity was associated with physical activity, a lifestyle factor reflecting resilience. However, this connectivity was not significantly related to other proxies of cognitive reserve (education and IQ)^9^, suggesting that lifestyle-driven resilience mechanisms may operate through distinct neural pathways. This is consistent with previous reports suggesting social factors such as education likely act on brain networks independent of those affected by Aβ^63^. Our results emphasize the need to incorporate such resilience variables when examining the relationships between AD pathology and brain function.

Physical activity (PA) is one potential modifiable factor for reducing the risk of cognitive decline and dementia^12,64^, however pathways underlying its resilience against developing dementia are unknown. PA has been associated with better cognitive function^65,66^, as well as protective effects against amyloid-related cognitive decline and neurodegeneration independent from vascular risk^11^. However, reports on associations between PA and Aβ are often mixed^67,68^. For example, studies showed PA benefit cognition mediated by reduction in Aβ^69^, whereas some studies found the relationship is influenced by APOE ε4 status^70–72^, and evidence also suggest the associations are likely stronger in the very early phase of AD^73^. PA has also been related to tau pathology^74^. More recent studies have shown PA provides resilience against influence from tau on executive function, and no interaction with episodic memory and Aβ^75,76^. Consistent with these literatures, our results suggest that PA-related network pathways are associated with Aβ burden, which is one of the initiating pathological events in AD. However, these PA-related resilience pathways did not moderate the effect of Aβ on working memory performance, suggesting that they might preferentially support other non-memory cognitive domains such as executive function. Future studies are needed to test this possibility.

Our results provided evidence that Aβ might trigger network reconfiguration, where the altered DMN-DAN connectivity represents a pathway of physical activity on resilience. However, the connectivity did not significantly moderate the influence of amyloid on task performance, which is consistent with literature showing no interaction between PA and amyloid on cognition^75^. Previous studies have reported that PA is linked to stronger connectivity in the DMN, salience, and control networks^10,77–80^. PA has also been closely linked to couplings between the DMN and the FPN, where DAN is also named the dorsal FPN^81^. An intervention study showed that 12 weeks of aerobic exercise increased connectivity within posterior DMN regions, as well as between posterior DMN regions and the left postcentral gyrus of DAN in both cognitively normal and MCI older adults^82^. Another randomized controlled trial found that exercise training modulated connectivity within the FPN^83^. Increased DMN coupling with the FPN has been related to cognitive flexibility^84^, creativity in older adults^85^, and proposed as a sign of adaptive compensatory mechanism^86,87^. In cognitively normal individuals, greater DMN-DAN coupling has been associated with better memory performance over time, possibly reflecting compensatory mechanisms, whereas higher DMN-DAN coupling relates to worse performance in MCI^56^. This stage-dependent effect suggests a temporally compensatory resilience mechanism but eventually relates to network failure as pathology progresses^1^. In a preliminary longitudinal analysis, we observed that baseline PA predicted changes in DMN-DAN connectivity over time. This result supports the idea that higher DMN-DAN coupling might serve as a marker of resilience in cognitively normal older adults, and physical activity appears to facilitate such adaptive network integration in the face of Aβ pathology.

Studying associations between early AD pathology and brain function, as well as their relationships to task performance and resilience factors, have important implications for developing targeted intervention design and functional biomarkers for monitoring interventional effects. For example, given that amyloid-related connectivity changes were phase-specific (prominent during the probe/retrieval phase), future work could test phase-dependent modulation using interventions such as TMS to improve task performance. Prior studies have shown that the exact timing of brain stimulation relative to the phase during working memory tasks is critical and exhibits distinct effects on performance^88^. Similarly, exercise interventions have been found to modulate retrieval-related fMRI activation in MCI^30^. These findings highlight the potential value of brain functional measures for guiding and evaluating targeted interventions that promote resilience. Future studies should test whether interventions such as exercise or brain stimulation can modulate DMN-DAN connectivity in supporting working memory retrieval or improving cognition in aging.

When investigating the relationship between AD pathology and brain function, most prior studies have focused on correlation-based resting-state functional connectivity or task-evoked brain activation. However, recent work has demonstrated that task-modulated connectivity captures context-dependent brain reconfiguration, which cannot be identified using correlation-based analyses of spontaneous fluctuations or task co-activations^17^. Only a limited number of studies have examined task-modulated connectivity in studies of AD pathology. Integrating task connectivity with AD biomarkers (amyloid, tau, cortical thickness) will be crucial for understanding the neuropathological progression of AD, as well as pathways linking pathology to cognition and modifiable resilience factors. For example, based on a task paradigm involving novel and repeated scenes and objects recognition, a recent study examined amyloid-related brain network disruptions using task connectivity via dynamic causal modeling (DCM)^18^. Their results showed that Aβ-related alterations in the DMN drive medial temporal lobe (MTL) hyperactivity and promote early tau accumulation in the entorhinal cortex, providing evidence on how Aβ-associated network-to-network interactions relate to Aβ-tau interaction. We used a computationally simpler but conceptually related approach, i.e. PPI analysis, which enables examination of whole-brain task-modulated connectivity rather than being limited to a few predefined ROIs or networks.

Characterizing connectivity at multiple spatial scales, ranging from brain regions to networks, might provide a more comprehensive view to help understand how Aβ pathology affects brain function. Recent studies highlight the importance of considering multiple spatial scales analysis when studying brain function^89^. Our analyses at both the nodal and network levels revealed distinct results, while regional node strength mediated amyloid effects on cognition, network-level connectivity did not, but demonstrated a significant association to physical activity. This might suggest amyloid-related local vulnerability of DMN regions, while network-level DMN-DAN couplings represent network adaptive resilience in the face of Aβ. Local disruptions in DMN hub regions may have more immediate behavioral consequences than broad network disconnections. Future studies should consider multiLscale connectivity analyses to study AD pathology relationships to brain function, such as cross-scale analyses or multi-scale network models^90^.

This study has several limitations that warrant consideration. First, tau pathology was not included due to limited sample availability, as tau-PET data were only available at follow-up and not at baseline in our study. Given that tau pathology often demonstrates a closer relationship with cognitive performance and physical activity, and that numerous studies showing the interactions between Aβ and tau^91,92^, future studies should incorporate tau-PET imaging.

However, a recent resting-state fMRI study suggests that DMN-DAN connectivity represents cognitive dysfunction independent of tau pathology^93^. The present study focused on Aβ pathology because it is considered one of the earliest events in the AD cascade, making it particularly relevant for understanding early brain functional changes in cognitively normal older adults. Additionally, in this study, we did not account for APOE ε4 status, which may influence the relationship between physical activity and Aβ burden. Second, physical activity was assessed using self-reported questionnaires, which are subject to potential recall bias and measurement error^94^. Future studies should consider objective approaches, such as accelerometry, to improve reliability. Third, the sample size was relatively small and our sample composed primarily of cognitively normal, amyloid-negative participants. Replication in larger cohorts and including individuals along the AD continuum such as those with MCI, will be important to confirm the generalizability of our findings. Future studies with longitudinal designs are needed to clarify the temporal sequence between Aβ accumulation and brain connectivity alterations. However, prior studies suggest that these relationships may be bidirectional, forming a “vicious cycle” in which pathological accumulation alters brain networks, which in turn may facilitate further pathological spread^95^. Finally, although our exploratory longitudinal analysis suggested that physical activity precedes changes in brain connectivity, interventional studies will be crucial to establish causal links between lifestyle resilience factors and brain function. The current findings may provide potential functional markers for monitoring lifestyle-based intervention outcomes, as well as for informing target selection in brain stimulation aimed at promoting resilience in aging.

## Conclusion

In summary, this study demonstrates that higher Aβ burden is associated with reduced task-modulated connectivity supporting working memory retrieval. Specifically, regional nodal connectivity, rather than system-level network coupling, mediated the relationship between Aβ burden and working-memory performance, suggesting the nodal vulnerability of posterior-medial and default mode hub regions to early AD pathology. Network-level couplings, especially between the default mode and dorsal attention networks, were associated with physical activity, rendering a potential lifestyle-related resilience pathway that supports brain function in the presence of early AD pathology. Together, these findings provide insight into early-stage brain functional vulnerability to Aβ burden and identify potential network targets that may inform monitoring and guide intervention strategies aimed at promoting resilience in aging.

## Data availability

Demographic information, PET measures, and brain connectivity data generated for this study are publicly available at https://github.com/hehengda/AmyloidLTS.git. Raw and preprocessed datasets analyzed during the current study are available from the corresponding author upon reasonable request.

## Funding

This work was supported by the National Institutes of Health/National Institute on Aging (NIH/NIA; grant numbers R01 AG038465 and R01 AG026158).

## Competing interests

The authors report no competing interests.

## Notes

### Competing Interest Statement

The authors have declared no competing interest.

