## Supplementary Material for "Amyloid-beta relates to default and attention network connectivity during working memory retrieval"

### **Supplementary text**

##### **Resting-state functional connectivity analysis**

For comparison purposes, correlation-based functional connectivity was also estimated from the resting-state fMRI dataset. Resting state fMRI scans were collected with a T2*-weighted echo-planar imaging (EPI) sequence (TR/TE = 2000/20 ms; flip angle = 72°; FOV = 224 × 224 mm; in-plane resolution = 112 × 112 voxels; slice thickness = 3 mm; 37 axial slices; either 150 or 285 volumes). Three dummy EPI volumes acquired at the beginning of each fMRI acquisition run were excluded before data preprocessing.

Resting-state fMRI data were preprocessed using an in-house pipeline [Razlighi et al., 2014]. Slice-timing correction was applied with Fourier‐space phase shifting (FSL v6.0.4)[Smith et al., 2004], followed by rigid‐body motion correction of all volumes with the first time point volume as the reference volume [Jenkinson et al., 2002]. A mean EPI reference image was generated, and framewise displacement (FWD) and root mean square difference (RMSD) were computed as motion parameters. Volumes with FWD > 0.5 mm or RMSD > 0.3% were scrubbed and replaced by linear interpolation of neighboring frames. Data were temporally filtered (0.01-0.08 Hz) using a nonlinear high-pass and Gaussian low-pass filter. After motion correction, scrubbing, and temporal filtering, nuisance regressors (FWD, RMSD, bilateral white-matter signals, and lateral ventricle signals) were removed. No global signal regression was performed. Mean time series were then extracted from 200 cortical Schaefer ROIs. Pairwise Pearson correlations were calculated and Fisher z-transformed to generate normalized functional connectivity matrices with zero diagonals. Quality control excluded participants with zero (or very low) variance in any ROI time series.

##### **Regional node strength mediates the association between amyloid-beta burden and task performance**

For the global standardized uptake value ratio (SUVR) measure, mediation analysis revealed a significant indirect effect of amyloid burden on task performance through regional node strength (mediation effect = -0.0811, 95% confidence interval = [-0.1741, -0.0166], p < 0.0064). In contrast to the signature-region analysis, the direct effect of global amyloid burden on task performance remained significant (c′ = -0.2353, 95% confidence interval = [-0.4315, -0.0182], p < 0.0344).

##### **Cross-validation analyses with different fold numbers**

Using the 20-fold cross-validation (CV) framework described in the main text, averaged regional node strength significantly predicted task performance (permutation p < 0.0020), while averaged network connectivity significantly predicted physical activity (permutation p < 0.0130). To evaluate the stability of these findings across different CV schemes, additional analyses were conducted using K = 5, 10, and 15 folds. Results were consistent for regional node strength in predicting task performance (K = 5: permutation p < 0.0020; K = 10: permutation p < 0.0020; K = 15: permutation p < 0.0020). For network-level connectivity, the results also remained consistent when predicting physical activity (K = 5: permutation p < 0.0100; K = 10: permutation p < 0.0100; K = 15: permutation p < 0.0080). Together, these results demonstrate that predictive associations were robust to the choice of fold number in the cross-validation procedure.

##### **Amyloid-related network-level connectivity is related to physical activity**

Average amyloid-related network-level connectivity was significantly associated with physical activity (r = 0.3931, p < 0.0011; Pearson correlation controlling for age and sex). However, these network-level connections did not significantly moderate the effect of amyloid burden on task performance. No significant moderation effect of connectivity on the relationship between amyloid burden and task performance was observed (testing on the interaction term: β = 0.0062, t = 0.2073, p = 0.8363).

| 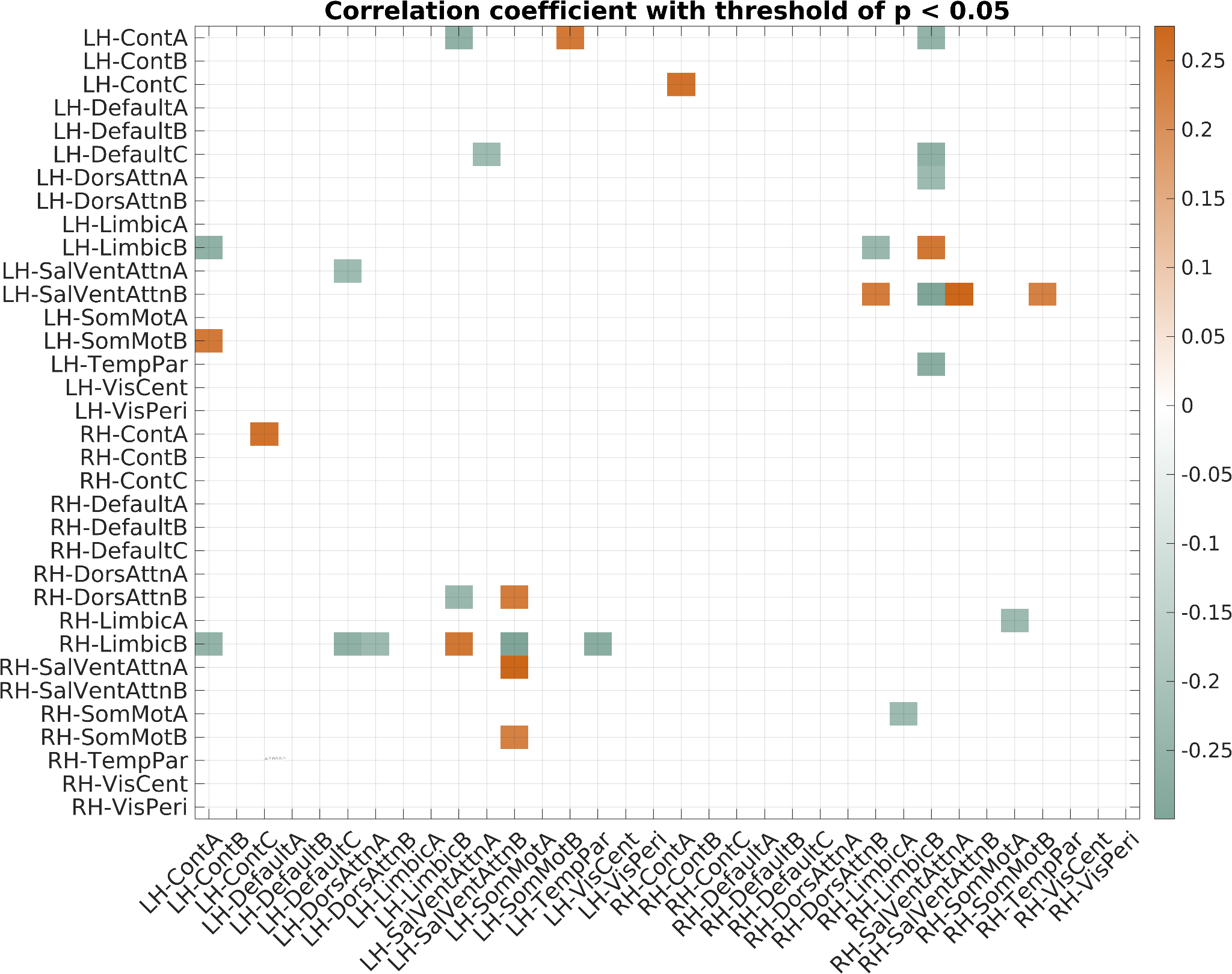 | 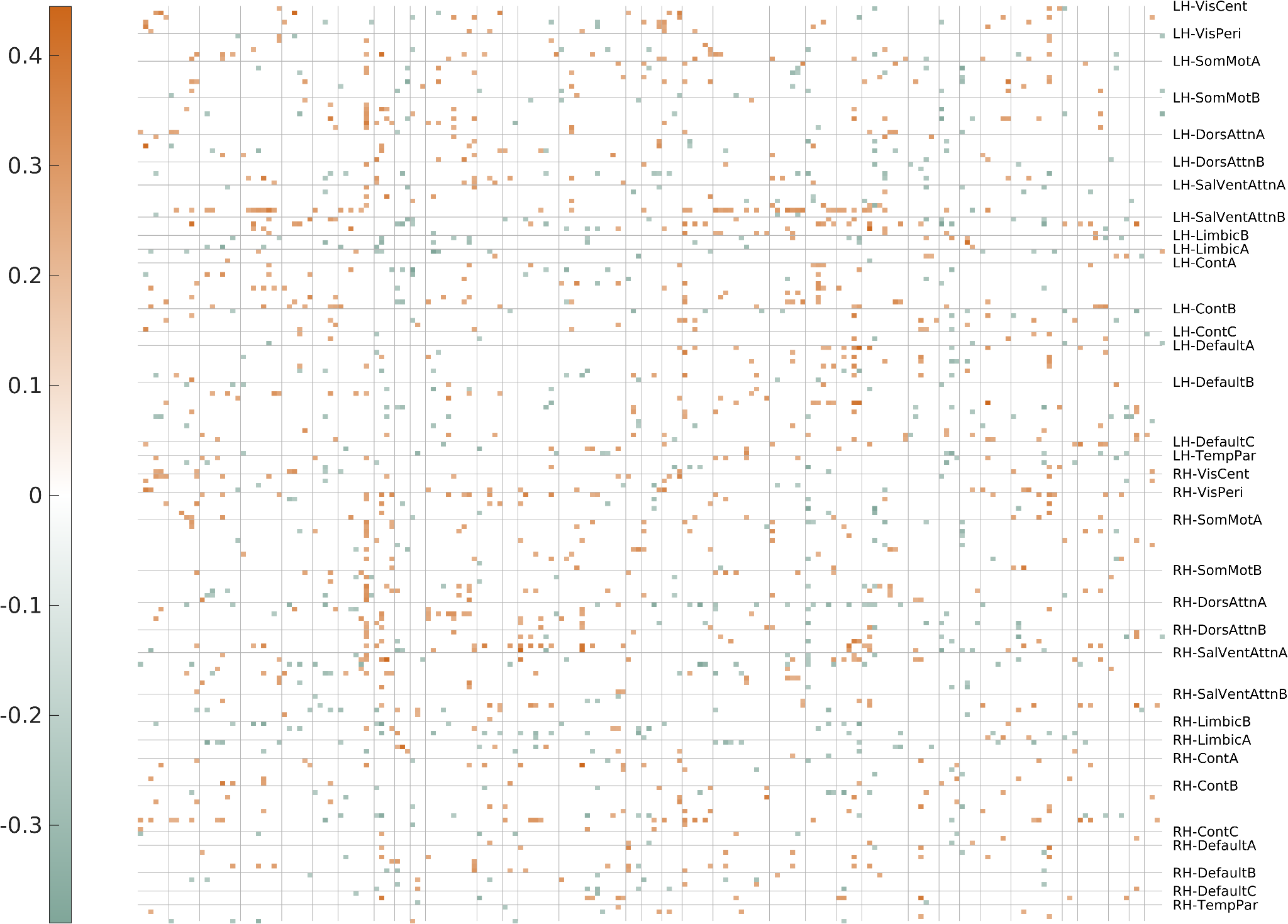 |
| --- | --- |

**Figure S1. Amyloid-associated changes in resting-state functional connectivity with uncorrected p < 0.05.** There is no significant correlation between amyloid burden in the signature regions and resting-state functional connectivity (false discovery rate correction q < 0.05). Correlation was estimated with partial Spearman correlation controlling age and sex. The relationships were tested both at the network-level and node-level.

**Table SI.** Associations between amyloid burden and node-level connectivity during the probe phase (SUVR in signature regions and global SUVR).

|  | SUVR in signature regions | | Global SUVR | |
| --- | --- | --- | --- | --- |
|  | r | p | r | p |
| Connection of right retrosplenial area and the right somatomotor area | *-0.5053* | *1.4980E-06 (**)* | *-0.4276* | *0.0001 (**)* |
| Node strength of LH_DefaultC_PHC_1 | *-0.4610* | *1.4859E-05 (**)* | *-0.2885* | *0.0090 (**)* |
| Node strength of RH_ContB_IPL_2 | *-0.3754* | *0.0006 (**)* | *-0.2189* | *0.0496 (*)* |
| Node strength of RH_DefaultC_Rsp_1 | *-0.3791* | *0.0005 (**)* | *-0.2652* | *0.0167 (*)* |
| Node strength of RH_DefaultC_PHC_1 | *-0.3999* | *0.0002 (**)* | *-0.2624* | *0.0179 (*)* |

** p < 0.05; ** p < 0.01*

**Table SII.** Associations between amyloid burden and network-level connectivity during the probe phase (SUVR in signature regions and global SUVR).

|  | SUVR in signature region | | Global SUVR | |
| --- | --- | --- | --- | --- |
|  | r | p | r | p |
| RH-DefaultC and LH-DorsAttnA | *-0.3918* | *0.0003 (**)* | *-0.2157* | *0.0531* |
| RH-DefaultC and LH-DorsAttnB | *-0.4884* | *3.7321E-06 (**)* | *-0.3472* | *0.0015 (**)* |
| RH-DefaultC and RH-DorsAttnB | *-0.4271* | *0.0001 (**)* | *-0.2195* | *0.0490 (*)* |
| RH-DefaultC and RH-SomMotA | *-0.4885* | *3.7214E-06 (**)* | *-0.3453* | *0.0016 (**)* |
| LH-DefaultC and RH-DorsAttnB | *-0.4071* | *0.0002 (**)* | *-0.3010* | *0.0063 (**)* |

** p < 0.05; ** p < 0.01*
